# The molecular mechanisms underlying hidden phenotypic variation among metallo-ß-lactamases

**DOI:** 10.1101/422303

**Authors:** Raymond D. Socha, Nobuhiko Tokuriki

## Abstract

Genetic variation among orthologous genes has been largely formed through neutral genetic drift to maintain the same functional role. In some circumstances, however, this genetic variation can create critical phenotypic variation, particularly when genes are transferred to a new host by horizontal gene transfer (HGT). Unveiling “hidden phenotypic variation” through HGT is especially important for genes that confer resistance to antibiotics, which continue to disseminate to new organisms through HGT. Despite this biomedical importance, our understanding of the molecular mechanisms that underlie hidden phenotypic variation remains limited. Here we sought to determine the extent of hidden phenotypic variation in the B1 metallo-β-lactamase (MBL) family, as well as to determine its molecular basis by systematically characterizing eight MBL orthologs when they are expressed in three different organisms (*E. coli, P. aeruginosa, and K. pneumoniae*). We found that these MBLs confer diverse levels of resistance in each organism, which cannot be explained by variation in catalytic efficiency alone; rather, it is the combination of the catalytic efficiency and abundance of functional periplasmic enzyme that best predicts the observed variation in resistance. The level of functional periplasmic expression varied dramatically between MBL orthologs and between hosts. This was the result changes at multiple levels of each enzyme’s functional: 1) the quantity of mRNA; 2) the amount of MBL expressed; and 3) the efficacy of functional enzyme translocation to the periplasm. Overall, we see that it is the interaction between each gene and the host’s underlying cellular processes (transcription, translation, and translocation) that determines MBL genetic incompatibility thorough HGT. These host-specific processes may constrain the effective spread and deployment of MBLs to certain host species, and could explain the current observed distribution bias.

**Author Summary:** Orthologous genes spread among different organisms, typically maintaining the same functional role within the cell while accumulating some, presumably functionally-inert, genetic variation over time. However, these seemingly neutral gene sequence changes among orthologs can be revealed to have substantial difference in protein phenotypes, and thus, organismal fitness, when they are transferred to other host species. This so-called “hidden phenotypic variation” through horizontal gene transfer may play an important role in dissemination of antibiotic resistance genes, in particular. In this work, we systematically investigated the extent of phenotypic variation in eight orthologous antibiotic resistant genes from the metallo-β-lactamases family (MBLs), and identified the molecular causes underlying the observed phenotypic variation. We found that functional protein expression varied substantially among MBLs (causing significant variation in the level of antibiotic resistance conferred), and that this could not be explained by variation in catalytic efficiency alone. Instead, we see that functional variation is caused by multiple steps in the protein production, transcription, translation and translocation, that are necessary to provide functional enzymes in the bacterial periplasm. Thus, the successful gene transfer and dissemination of antibiotic resistance genes can be determined by complex interactions between the gene and host underlying cellular processes.

## Introduction

Orthologs are genetically diverged genes that typically perform the same functional role in different host organisms. The genetic variation among orthologs is largely thought to be neutral, or not related to their function, and acquired through genetic drift, i.e., a process in which a purifying selection maintains physiological function (1). However, some genetic differences may be driven by adaptation resulting from, for example, transient historical changes in the level of selection pressure or in the cellular environment (e.g., pH, temperature, the concentration of nutrients and antibiotics) (2). Moreover, some genetic changes may be driven by co-evolution with other proteins and molecules within each organism (3). While these sequence changes can be neutral within each organism, they can be deleterious or advantageous if the gene is passed to a new host organism through horizontal gene transfer (HGT), revealing a “hidden phenotypic variation” (4-7). Such hidden phenotypic variation often manifests in the laboratory as sub-optimal heterologous protein expression in conventional laboratory hosts such as *Escherichia coli*, which occurs due to incompatibility between the gene and host (8-11). Moreover, hidden phenotypic variation may play an important role in the evolution and dissemination of antibiotic resistance genes because they are frequently transferred between organisms through HGT (4,5,12). However, our understanding of the molecular mechanisms underlying phenotypic variation and gene incompatibility is still limited.

The B1 Metallo-β-Lactamases (MBL) family is one such group of antibiotic resistance genes that has disseminated extensively through HGT to a wide variety of bacterial pathogens, causing a serious threat to healthcare systems worldwide (13-16). Despite high levels of genetic variation (with pairwise amino acid identities as low as 20%), these orthologs feature a shared αββα-fold, an identical active-site architecture (two zinc binding sites coordinated by H-H-H and D-C-H), and the ability to confer resistance to most β-lactam antibiotics by catalyzing the hydrolysis of the β-lactam ring (**Figure 1**) (17-19). Since the first B1 MBL, BcII, was isolated from the genome of *Bacillus cereus* in 1966, MBLs have been identified in a growing number of bacteria that belong to diverse phyla such as Bacteroidetes, Firmicutes and Proteobacteria (20-24). However, in the last two decades, some types of MBLs, such as NDM-type, VIM-type, IMP-type, and SPM-type have become “acquirable” because of their association with plasmids and other transferable elements, and have subsequently disseminated to diverse clinical pathogens, such as *Pseudomonas aeruginosa, Acinetobacter baumanni, Klebsiella pneumoniae, Enterobacter cloacae,* and *Escherichia coli* (18,25-28). These observations lead to a series of questions regarding phenotypic variation among MBLs and antibiotic resistance in pathogens: How might phenotypic variation manifest within this family? Are there phenotypic differences between the acquirable and chromosomally encoded MBLs? Moreover, what are the molecular properties determine phenotypic variation? To date, there has been no study that systematically compares the MBLs and their diverse hosts organisms to address these questions. Understanding the genetic and molecular causes underlying phenotypic variation among MBLs is important in developing our ability to prevent and control the dissemination of multi-drug resistance genes.

**Figure 1:**
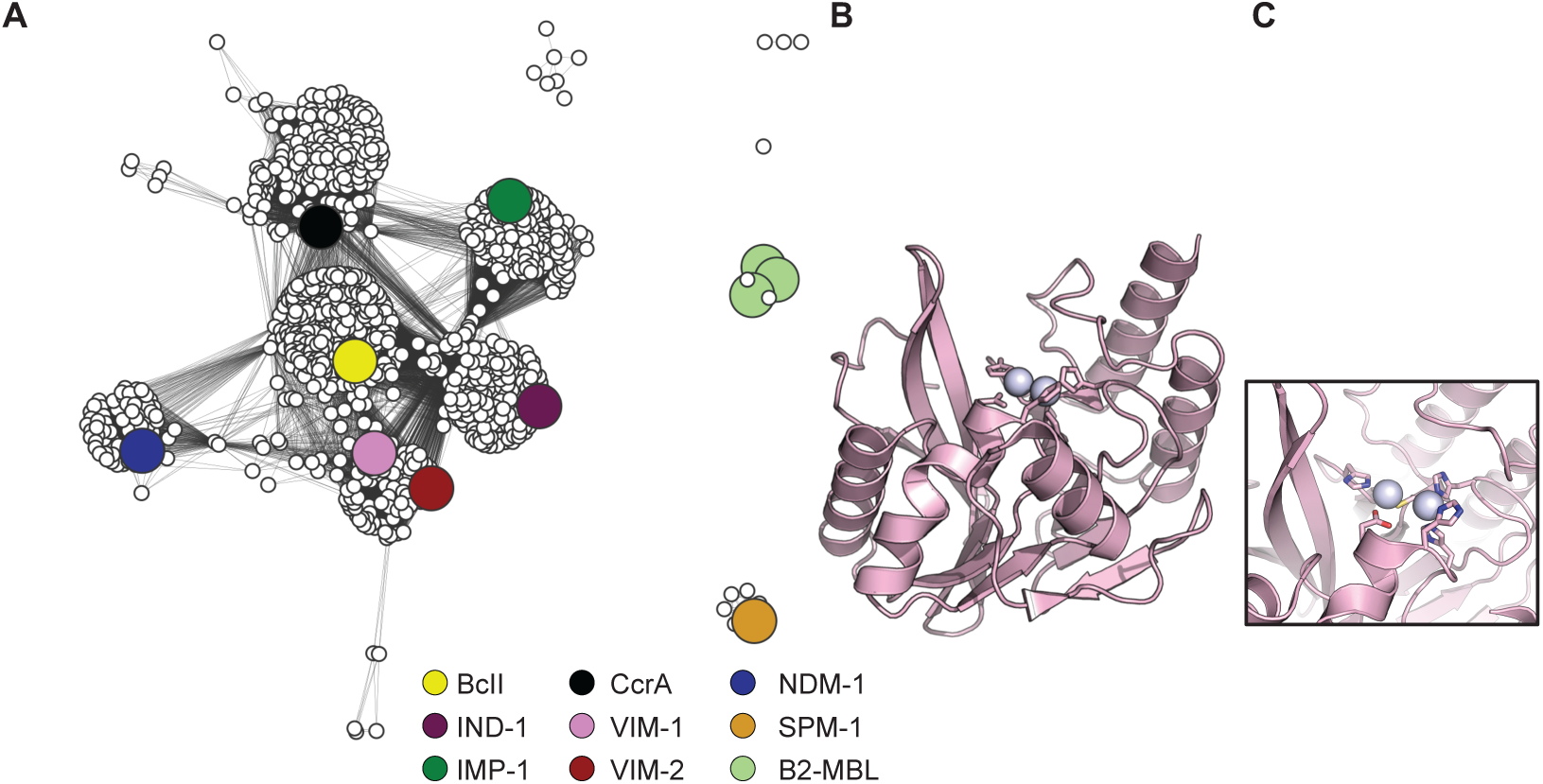
Sequence similarity networks for the metallo-β-lactamase family, with a representative crystal structure. **A.** The sequence similarity network within the B1 and B2 MBL family with 1224 sequences visualized with a BLAST *e*-value cutoff of 1e^-55^. The sequence of the MBLs are used in the study was shown as large circles and highlighted with colours: BcII (yellow), IND-1 (purple), CcrA (black), NDM-1 (blue), VIM-1 (pink), VIM-2 (red), IMP-1 (green), SPM-1 (orange), and the B2 MBL family (light green). Sequence identities between the MBLs are presented in **Table 1** and **Supplementary Table 1**. **B.** Cartoon presentation of the crystal structure of VIM-2 (PDB ID: 1K03). **C.** The close-up view of the VIM-2 MBL active site that is conserved throughout the entire family: two zinc ions (shown as grey spheres) held in place by metal binding residues H-H-H and D-C-H (shown as sticks).

**Table 1.** Information of the eight MBLs that are used in this study.

|  | Amino acid identity (%) |  |  |  |  |  |  |  | Mobility<sup>1</sup> | Common host organisms<sup>2</sup> | Accession number<sup>3</sup> |
| --- | --- | --- | --- | --- | --- | --- | --- | --- | --- | --- | --- |
|  | BclI | IND-1 | IMP-1 | CcrA | VIM-1 | VIM-2 | NDM-1 | SPM-1 |  |  |  |
| **BclI** | 100% |  |  |  |  |  |  |  | Chromosomal | *Bacillus cereus* | AAA22276 |
| **IND-1** | 33% | 100% |  |  |  |  |  |  | Chromosomal | *Chryseobacterium indologenes* | ABO21411 |
| **IMP-1** | 33% | 30% | 100% |  |  |  |  |  | Acquired | *Pseudomonas aeruginosa*,<br>*Acinetobacter baumannii*, *Serratia marcescens*, *Enterobacter cloacae* | AAN87168 |
| **CcrA** | 31% | 28% | 33% | 100% |  |  |  |  | Chromosomal | *Bacillus fragilis* | AAA22904 |
| **VIM-1** | 36% | 27% | 30% | 29% | 100% |  |  |  | Acquired | *Pseudomonas aeruginosa*,<br>*Klebsiella pneumoniae*, *Serratia marcescens* | CAE46717 |
| **VIM-2** | 36% | 25% | 32% | 28% | 90% | 100% |  |  | Acquired | *Escherichia coli* | AAK26253 |
| **NDM-1** | 28% | 23% | 31% | 29% | 35% | 34% | 100% |  | Acquired | *Escherichia coli*, *Klebsiella pneumoniae*, *Enterobacter cloacae*,<br>*Acinetobacter baumannii* | AFI72857 |
| **SPM-1** | 26% | 29% | 32% | 24% | 26% | 26% | 23% | 100% | Acquired | *Pseudomonas aeruginosa* | AAR15341 |

1. Mobility indicates if the MBL is identified as a chromosomal copy in a single spece of bacteria or the MBL has been transferred to multiple bacteria (acquired).
2. Common host organisms indicate the original host of the MBL or the hosts that acquired the MBL and their closely related homologous (e.g., NDM-type). The information was obtained from Reference 27.
3. Accession number indicates National Center for Biotechnology Information (NCBI) accession no.

Here, we conduct a comprehensive characterization of eight orthologous MBL enzymes to determine their resistance level in three bacterial hosts, and unveil the extent to which genetic variation causes hidden phenotypic variation among them. We further performed diverse biochemical and biophysical characterizations of the enzymes to reveal the underlying molecular basis for the observed phenotypic variation.

## Results

### Metallo-β-lactamases provide diverse levels of antibiotic resistance

We chose a set of eight MBLs to investigate the phenotypic variation in the B1-MBL family **(Figure 1** and **Table 1)**: Three enzymes represent chromosomally-encoded MBLs (BcII from *Bacillus cereus*, IND-1 from *Chryseobacterium indologenes*, and CcrA from Bacteroides fragilis), and five represent those acquired on mobile genetic elements and have been identified in multiple bacterial pathogens (NDM-1, VIM-1, VIM-2, IMP-1, SPM-1). These enzymes also reflect the high sequence diversity within the B1-MBL family (amino acid identities between the enzymes range from 23% to 36%, except between VIM-1 and VIM-2, which is 90%) (**Figure 1**, **Table 1** and **Supplementary Table 1**). The MBL genes were subcloned into a modified broad-host-range pBBR1MCS-2 vector (29) along with the inducible P_BAD_ promoter (pBBR1-pBAD) and subsequently transformed into three bacterial strains: *Escherichia coli* E. cloni^®^ *10G, Pseudomonas aeruginosa* PA01, and *Klebsiella pneumoniae* ATCC13883. These three organisms were chosen because they represent major opportunistic pathogens that have recently acquired MBLs through HGT. The minimum inhibitory concentration (MIC) for the strains harbouring the MBL genes was determined for six different compounds representing the three major classes of β-lactam antibiotics: cephalosporins, cefotaxime (CTX) and ceftazidime (CAZ); carbapenems, meropenem (MEM) and imipenem (IMI); and penams, ampicillin (AMP) and penicillin (PEN). The bacterial growth was measured on agar plates supplemented with each antibiotic (the range of concentrations screened for each antibiotic was: cephalosporins, 0.032 μg/mL to 4096 μg/mL; carbapenems, 0.016 μg/mL to 64 μg/mL; penams, 2 μg/mL to 32768 μg/mL). It should be noted that the endogenous MIC of each strain for each antibiotic varies considerably due to their intrinsic resistance; yet, we are able to quantify the contribution of MBLs to increase MICs in each strain (**Figure 2**).

**Figure 2:**
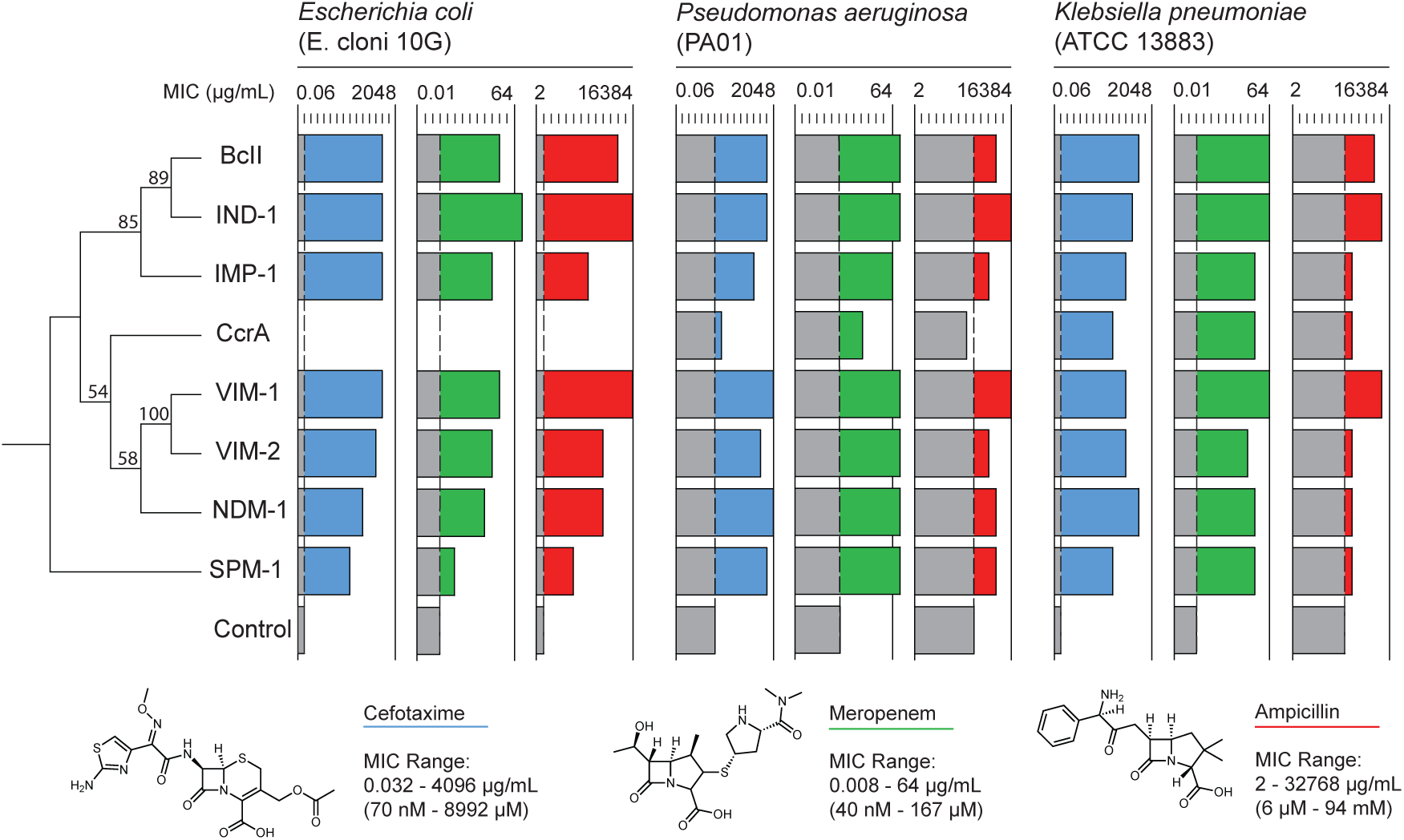
Measured minimum inhibitory concentrations (MICs) for the metallo-β-lactamases with representative β-lactam antibiotics in *E. coli, P. aeruginosa, and K. pneumoniae*. MICs for cefotaxime (blue), meropenem (green), and ampicillin (red) measured for each MBL (BcII, IND-1, IMP-1, CcrA, VIM-1, VIM-2, NDM-1, and SPM-1) in *E. coli* 10G, *P. aeruginosa* PA01, and *K. pneumoniae* ATCC 13883. MICs were determined with the concentration at which at least three of four replicates did not grow. The grey bar represents the background resistance level of the organisms without MBL expression. The chemical structures of cefotaxime, meropenem, and ampicillin as well as the concentrations screened to determine the MICs are shown below. MICs for ceftazidime, imipenem, and penicillin are presented in **Supplementary Figure 1.**

The MBLs confer diverse levels of resistance (**Figure 2**, **Supplementary Figure 1**, and **Supplementary Table 2**). For example, there is an over 4000-fold range in the ceftazidime MIC between MBLs in *P. aeruginosa*, from 2 μg/mL for CcrA to 8192 μg/mL for NDM-1. Nonetheless, within each organism, the relative order of the MBLs by resistance was similar for the six different antibiotics. For example, IND-1 and VIM-1 in *E. coli* generally confer the highest MICs, followed by BcII, VIM-2, NDM-1, IMP-1, SPM-1, and lastly, CcrA. This indicates that the all MBLs exhibit similar broad substrate specificity, but the level of resistance that the enzymes confer varies significantly. By contrast, a different trend emerges when comparing the resistance levels across the three different organisms. For example, SPM-1, which confers one of the lowest MICs among the eight MBLs in *E. coli* (*e.g.,* 32-fold lower ampicillin MIC than BcII, NDM-1, VIM-1 and IMP-1), provides a level of resistance equivalent to other MBLs such as BcII and NDM-1, and greater than IMP-1 in *P. aeruginosa*. CcrA, whose expression appears to be lethal in *E. coli* and thus did not result in formation of a colony even in the absence of antibiotics, is well tolerated in the other two organisms and even confers substantial resistance to *K. pneumoniae*. Taken together, the results suggest that the relationship between the MBL genes and their host organisms plays a strong role in determining the relative resistance that they provide.

### Variation in enzyme catalytic efficiency fails to explain the variation in antibiotic resistance

The level of antibiotic resistance is generally considered to be strongly associated with the associate antibiotic resistance enzyme’s catalytic efficiency (*k*_cat_/*K*_M_), and as a result it is a common measurement that is made upon the discovery of a new MBL variants (30-32). We examined to what extent the *k*_cat_/*K*_M_ of the MBLs can explain the variation observed in their respective MICs. The mature MBL genes (with their signal peptides removed) were fused with a C-terminal strep-tag by subcloning into the pET-26(b) vector. The fusion proteins were subsequently overexpressed in *E. coli* BL21 (DE3), purified, and their kinetic parameters (*k*_cat_, *K*_M_, and *k*_cat_/*K*_M_) were determined for the six β-lactam substrates, in addition to a generic substrate, CENTA (**Supplementary Figure 2** and **Supplementary Table 3**). Overall, the all MBLs are highly efficient enzymes for all seven substrates, with *k*_cat_/*K*_M_ values ranging from 10^4^ to 10^7^ M^-1^s^-1^. For each substrate, the variation in *k*_cat_/*K*_M_ between the MBLs is relatively small, with the difference between enzymes typically less than one order of magnitude (**Supplementary Table 3**). Interestingly, the relative enzyme catalytic efficiency rarely aligns with the level of resistance that the enzyme confers (**Figure 3**, and **Supplementary Figure 3**). Of the six antibiotics assessed with the three different organisms (18 combinations), the relationship between *k*_cat_/*K*_M_ and MIC was inversely correlated in 10 of the 18 cases. The 8 remaining cases showed a positive, but very weak correlation. This is highlighted in *E. coli* in particular, where SPM-1 exhibits a 10-fold higher *k*_cat_/*K*_M_ for ampicillin compared to VIM-2 (2.7 × 10^6^ *vs.* 2.7 × 10^5^ M^-1^s^-1^), yet SPM-1 confers 256-fold lower resistance against ampicillin compared to VIM-2 (64 *vs*. 16,384 μg/mL). Thus, whereas the existence of β-lactamase activity is essential to confer resistance to bacteria, the level of resistance that bacteria obtains cannot be explained by the kinetic parameters (*k*_cat_/*K*_M_), and must be strongly affected by other factors that are associated with the relationship between the host and the enzyme.

**Figure 3:**
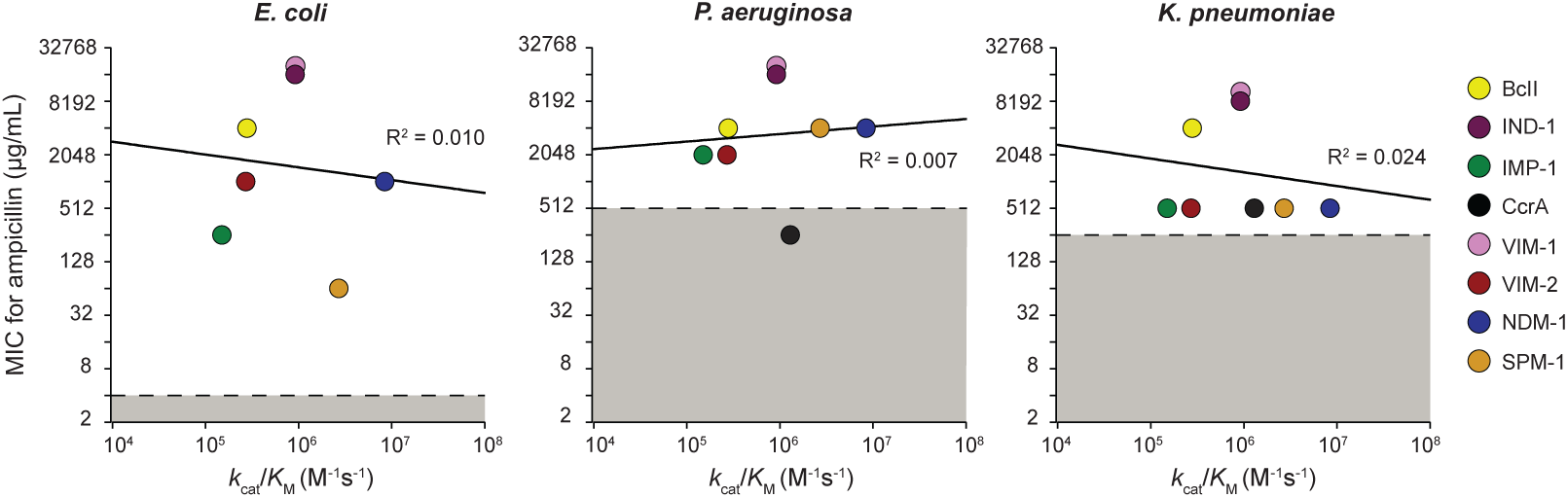
Relationship between *k*_cat_/*K*_M_ and ampicillin MICs for the MBLs in the three organisms. The measured ampicillin MIC values for each MBL in relation to their *k*_cat_/*K*_M_ for each organism are shown. The background resistance for each organism is presented as the grey box. The relationship for the other 5 antibiotics is shown in **Supplementary Figure 3**.

### The combination of catalytic efficiency and periplasmic expression of metallo-β-lactamases determines the level of antibiotic resistance

The level of antibiotic resistance, or the fitness of the host strain (*W*), that is conferred by an MBL gene is the result not only of their catalytic efficiency (*k*_cat_/*K*_M_), but also the abundance of functional enzyme in the periplasmic fraction of the cell ([*E*_p_]). With a simplistic view, this relationship can be defined as *W* ∝ *k*_cat_/*K*_M_ × [*E*_p_] (33,34). To determine if this relationship can sufficiently explain the variation in observed resistance (MIC), we determined the [*E*_p_] of each enzyme by measuring the level of enzymatic activity in the periplasm fraction. Briefly, the cells that expressed MBLs were harvested, the periplasmic fractions were isolated using the osmotic shock method, and the level of β-lactamase activity of in the periplasmic fraction was determined using 50 μM of CENTA. The number of functional MBL enzymes per cell in the periplasm fraction [*E*_p_] was then calculated using the kinetic parameters and the cell density of the cultures (see methods). It should be noted that two enzymes, NDM-1 and IND-1, showed very low activity in the periplasm fraction, and thus we omitted these enzymes from the following analyses. As a previous study demonstrated, NDM-1 is not a soluble periplasmic enzyme but rather localizes to outer membrane vesicles, and exhibits low concentrations in the periplasm despite consistently providing high levels of resistance (35,36). IND-1 also exhibited unexpectedly low level of activities in the lysate enzymatic assay, and we speculate that IND-1 may be unstable in our assay buffer.

Of the six remaining MBLs, there was significant variation in the calculated [*E*_p_]. The relative range in the variation of [*E*_p_] (over 1000-fold) was greater than that of *k*_cat_/*K*_M_ (less than 100-fold) (**Figure 4A** and **Supplementary Figure 4**). For example, the [*E*_p_] of MBLs differs by more than >600-fold in *E. coli,* with SPM-1 expressing <20 molecules per cell and VIM-1 being produced >8000 molecules per cell. The relative order for the [*E*_p_] of six MBLs is similar across the three organisms (**Supplementary Table 4**); VIM-1 is consistently the most highly expressed enzyme with approximately 10-fold higher abundance than the other MBLs by across all three organisms. By contrast, SPM-1 and CcrA are the lowest expressed enzymes in the three organisms. Nonetheless, the specific level of [*E*_p_] of each enzyme varies depending on the host organism (**Supplementary Table 4**). For example, VIM-2’s periplasmic expression in *P. aeruginosa* is tripled compared to *E. coli*, whereas BcII’s expression is halved. Therefore, the expression of each MBL enzyme is highly dependent on both its sequences and the host organism.

**Figure 4:**
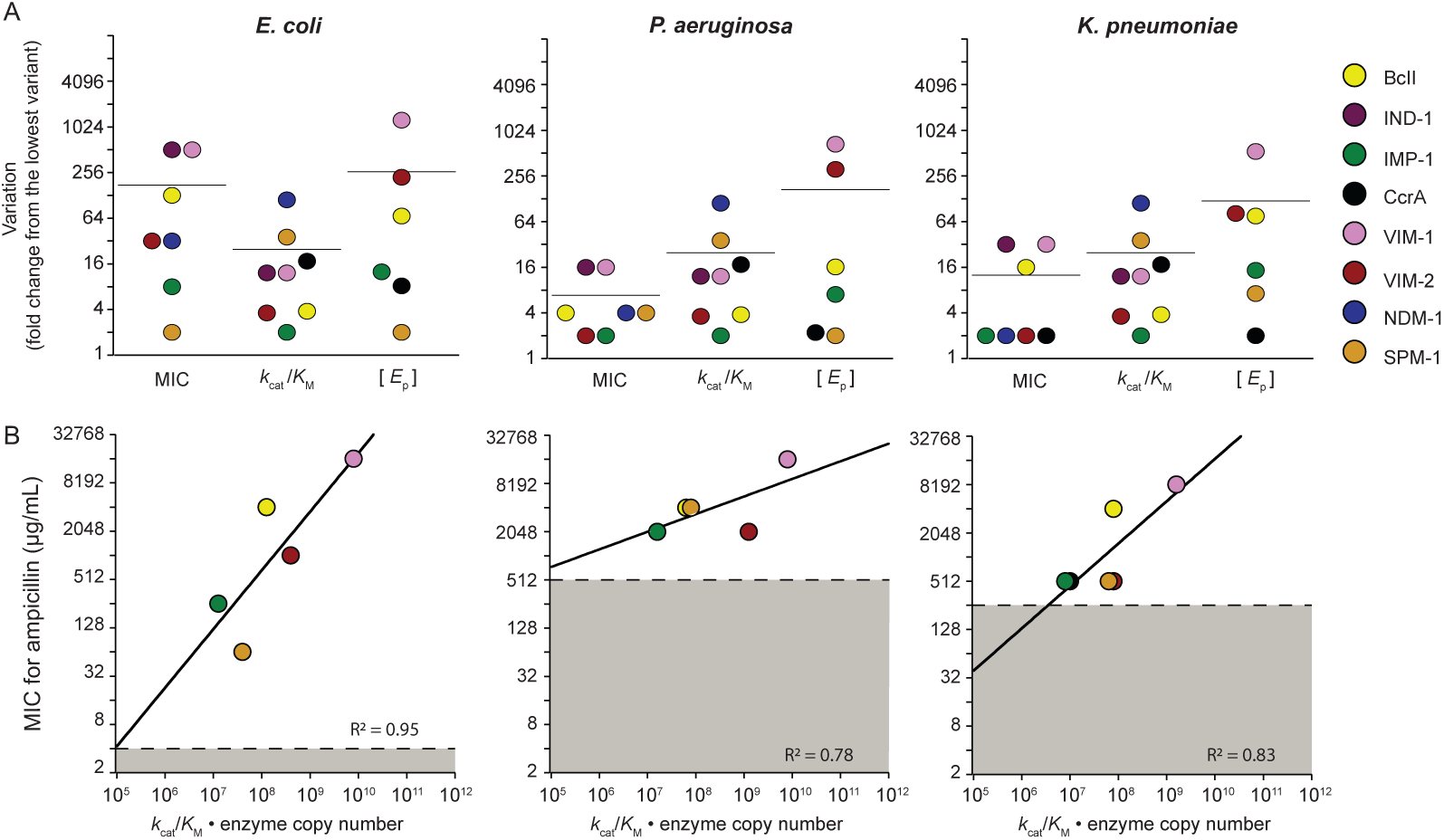
Relationship between MIC, *k*_cat_/*K*_M_, and [*E*_p_] for each MBL in the three organisms. **A.** The variation within MIC and the *k*_cat_/*K*_M_ for ampicillin, and the [*E*_*p*_] (the protein expression level in the periplasm fraction) within each organism for the MBLs. Each data point represents the fold-difference between the values for each MBL divided by the lowest value present for each property. The relationship for the other 5 antibiotics is shown in **Supplementary Figure 4**. **B.** The correlation between the product of *k*_cat_/*K*_M_, and [*E*_p_] with MIC. NDM-1 and IND-1 were not included in the fit as NDM-1 is a known to be bound to the outer membrane and could therefore not be accurately measured with the assay, while IND-1 activity was not detectable in the assay. The relationship for the other 5 antibiotics is shown in **Supplementary Figure 5**.

Overall, the variation in the antibiotic resistance levels conferred by MBLs is well-explained using the equation, log(*W*) = log(*k*_cat_/*K*_M_ × [*E*_p_]) (**Figure 4B** and **Supplementary Figure 4**), which captures the substantial variation in *k*_cat_/*K*_M_ and [*E*_p_] for each enzyme in each organism. The results suggest that each MBL’s MIC can be explained by variation in *k*_cat_/*K*_M_ and [*E*_p_]; however, it is noteworthy that MBL genetic variation appears to have a greater effect on expression rather than catalytic efficiency, suggesting it is the dominant mechanism driving variation in resistance conferred.

### The variation in MBL periplasmic expression reflects variation that emerges multiple processes of protein expression

Next, we sought to understand the underlying molecular mechanisms that determine MBL periplasmic expression. We determined the extent of variation in each factor that is associated with the overall variation in enzyme expression in the periplasm, e.g., the plasmid copy number, mRNA, and whole-cell enzyme expression, as well as the previously measured periplasmic protein concentration in *E. coli*. The variation between each MBL in plasmid copy-number and mRNA was measured by using quantitative PCR (qPCR). The enzyme abundance in the cytoplasm fraction ([*E*_c_]) was determined by lysing the spheroplasts (previously obtained during the periplasmic fraction isolation) and measuring the CENTA activity; the whole cell enzyme abundance ([*E*_w_]) is thus calculated by adding [*E*_c_] and [*E*_p_]. There is substantial variation in each molecule (DNA, RNA, and protein) among the eight MBLs (**Figure 5**): The variation is presented as normalized numbers that is relative to the lowest variant for each assay. As expected, the variation in the amount of the plasmid copy number was relatively small: only 3-fold difference between the MBLs. However, variation in the level of mRNA is notably larger, exhibiting up to 16-fold difference between MBLs, indicating that the processivity and/or stability of mRNA differ depending on the MBL sequence. Moreover, the range of variation between the eight MBLs further increased after protein expression, from 210-fold at the whole cell protein level, to 650-fold in the periplasmic protein level (**Figure 5A**). Interestingly, each enzyme behaves differently in each protein production process. For example, BcII exhibits the highest mRNA/DNA ratio, indicating that BcII mRNA is highly transcribed and/or stable in the cell (**Figure 5B-C**). On the other hand, VIM-1 exhibits a high protein/mRNA ratio suggesting that the high resistance of VIM1 is supported by high protein stability and/or translation rate (**Figure 5B-C**). Moreover, the fraction of translocated enzymes (the ratio between periplasm and whole protein) varies significantly depending on the gene sequence: VIM-1 and VIM-2 expressed more than 30% proteins in the periplasm, but SPM-1, IMP-1 and BcII showed only less than 10% of proteins are translocated (**Figure 5D**). This indicates that translocation is also a key factor in determining the MIC.

**Figure 5:**
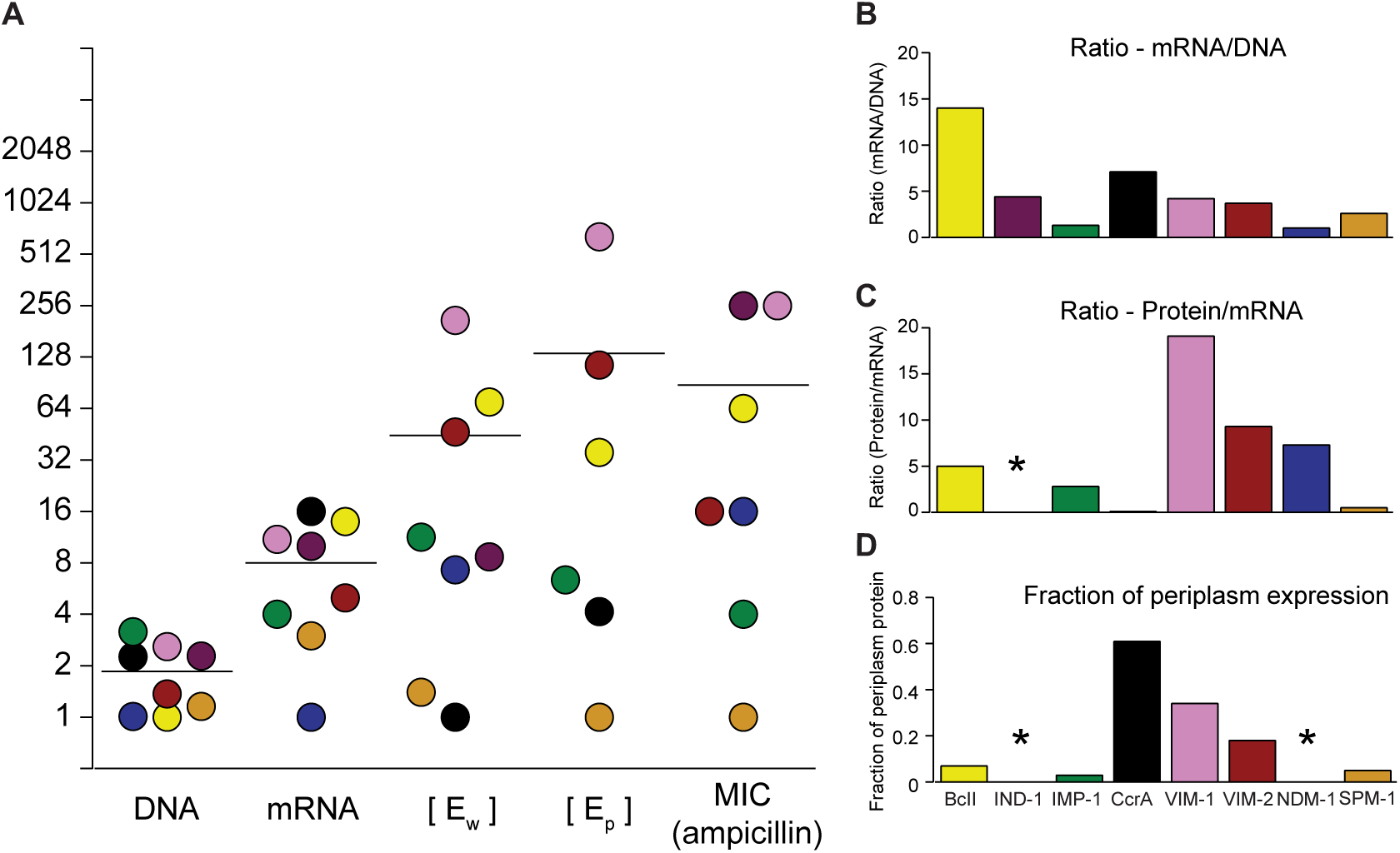
The variation between MBLs at each level of expression compared to the variation in ampicillin resistance.

Subsequently, in order to examine if the protein stability is correlated to the protein/mRNA ratio, we measured thermostability of the purified proteins. Interestingly however, we found that thermostability cannot explain variation in protein expression (**Supplementary Figure 5**). For example, VIM-1, and VIM-2 and NDM-1 exhibit comparatively high levels of protein expression, but they show relatively moderate thermostability (<60°C). Additionally, SPM-1 showed highest thermostability (73°C) despite exhibiting one of the lowest protein expression levels (**Supplementary Figure 5**). This indicates that more complicated properties of protein stability such as protein folding as well as translation processivity likely account for variation in protein expression.

Taken together, these results suggest that multiple properties such as mRNA, protein expression, stability and translocation underlie the phenotypic variation across the MBLs.

### A universal signal peptide does not fully abrogate phenotypic diversity

Establishing that translation and translocation are key factors in phenotypic variation, we sought to determine the effect of the signal peptide sequence on the MIC variation observed among MBLs. It has been demonstrated that the N-terminal region is strongly associated with the level of protein expression through its influence on translation initiation (37). In the case of the MBLs, this region also corresponds to the signal peptide of the translated protein, which would be expected to also exert significant influence on the efficacy of the MBL’s translocation to the periplasm (or outer membrane for NDM-type MBLs) through its interactions with the signal recognition particle, trigger factor, or SecA, in addition to the SecYEG translocase (38,39)}. This region also exhibits the highest level of genetic variation between the MBL genes, with no fully conserved residues and diverse lengths between 27 and 43 residues (as defined by where the first secondary structure element begins). Thus, we hypothesized that substitution of the native signal peptide of each MBL with a universal sequence may reduce the observed phenotypic variation.

The PelB leader sequence is a 22 amino acid signal peptide sequence that was originally identified from the pectate lyase B gene from *Erwinia carotovor* and is extensively used for recombinant periplasmic protein expression in *E. coli* (40). We replaced the native signal peptide of each MBL with the PelB sequence, generating PelB-MBL fusion genes in the pBBR1-pBAD vector, and determined the MICs of the three organisms harbouring the constructs for the six β-lactam antibiotics (**Supplementary Figure 6**). As predicted, the replacement of the signal peptide altered the MIC values; in some cases, the PelB peptide increased the MIC, for example causing a 2 to 8-fold increase for NDM-1 and SPM-1 in *E. coli.* However, for other MBLs, the substitution of the native signal peptides with PelB either led to no change or a decrease in MIC (**Figure 6A**). Consequently, the correlation between the PelB-MBL MICs and the measured catalytic efficiency became stronger compared to the MICs of the native MBLs (**Figure 6B** and **Supplementary Figure 6**). However, the correlations are still not as strong as those that take into account both catalytic efficiency and functional periplasmic expression. This suggests that while the sequences in the signal peptide region substantially contribute to the expression of the MBLs, and thus the antibiotic resistance of the host organisms, the phenotypic variation observed amongst the family cannot by explained solely by the signal peptide sequences alone. Genetic variation throughout the entire gene plays a role in determining the level of functional periplasmic expression.

**Figure 6:**
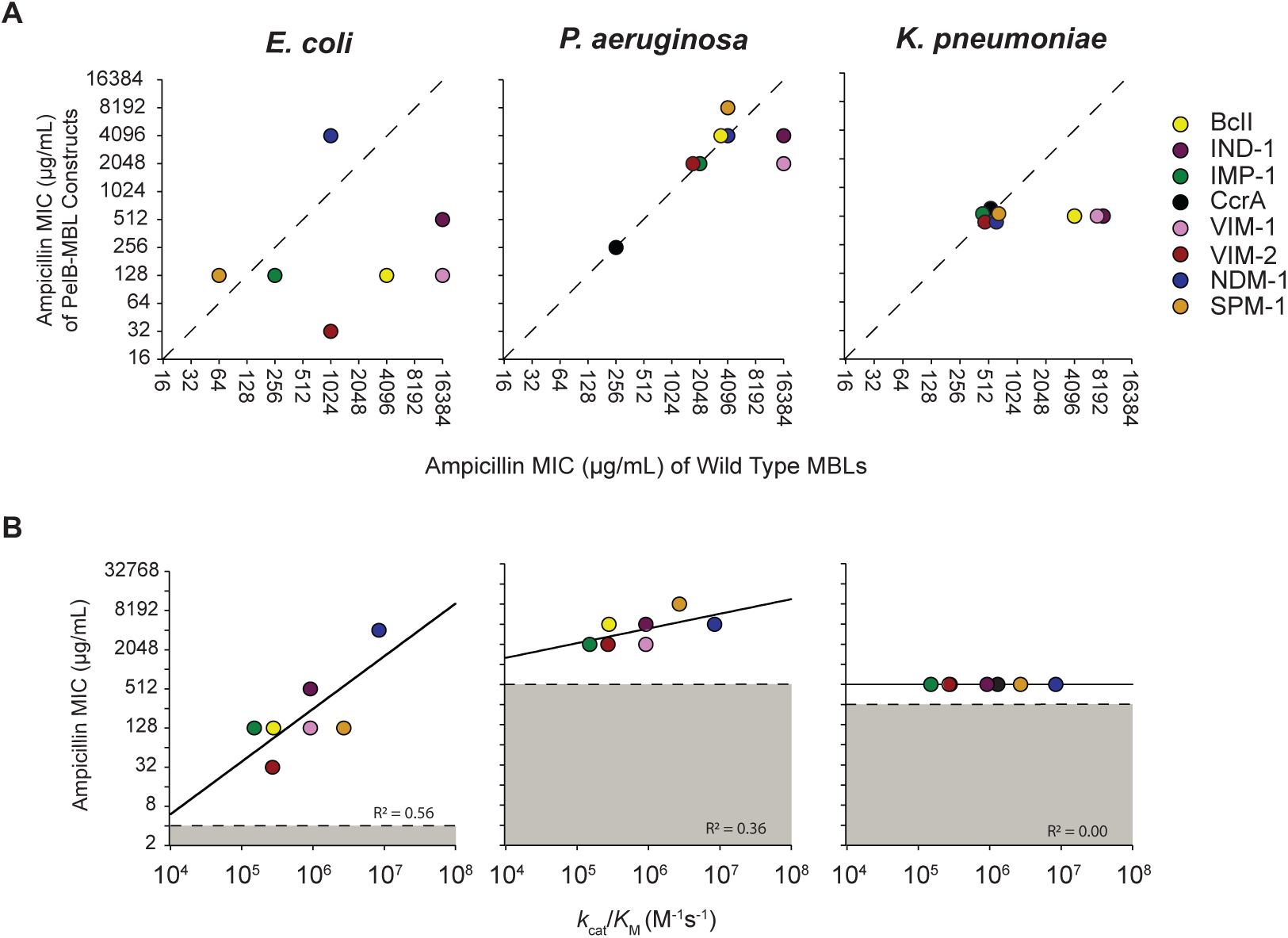
The effects of the replacement of the native signal peptide with the PelB leader sequence on MBL-conferred resistance. **(a)** Relationship between the MICs for each metallo-β-lactamase with their native signal peptide and with the PelB leader sequence for ampicillin, cefotaxime, and meropenem in *E. coli, P. aeruginosa,* and *K. pneumoniae*. **(b)** Correlation between the catalytic efficiency of each PelB-MBL (*k*_cat_/*K*_M_) and their ampicillin MICs. The relationship for the other 5 antibiotics is shown in **Supplementary**

## Discussion

With this work, we have systematically characterized eight B1-MBL enzymes and demonstrated that their genetic variation can cause substantial phenotypic variation upon expression in the same host organisms. Moreover, we see that strong variation is observed not only for enzyme catalytic efficiency but also in terms of the each enzyme’s functional periplasmic expression. Our study holds important implications for the genetic compatibility of genes that are horizontally transferred to new host organisms and the relationship between sequence and gene expression. What are molecular properties that determine protein expression and genetic compatibility? Many factors have been proposed such as GC content, codon usage, and mRNA folding energy, however, the evidence from independent studies is often contradictory (11,37,41-45). Indeed, a recent study that assessed 200 diverse antibiotic resistance genes in *E. coli* demonstrated that many proposed factors mentioned above cannot sufficiently explain the compatibility of the heterologous genes. Instead, only phylogenetic distance (*i.e.*, evolutionary distance between the original and new hosts) and mechanistic compatibility with the host (*i.e.*, dependency upon specific components of the host’s physiology and metabolism) provided a generally accurate estimation of protein fitness (5). Our observations suggest that variation in MBL fitness is caused by every step of the protein expression process, including transcription, translation as well as translocation. Thus, it is unlikely that a single parameter can be measured that explains genetic compatibility and protein expression in the cell, and it is essential to develop more comprehensive and multiple parameter models to understand genetic incompatibility.

Another important finding here is the observation that translocation is one of the mechanisms for gene compatibility. To date, a general trend in the MBL signal peptide sequence has been identified for each translocation system (e.g., the Sec protein translocation pathway) (46-50), however, we have little understanding of the relationship between the signal peptide sequence and its translocation efficacy, let alone the incompatibility between the translocation system of a particular organism and the signal peptide sequence of a particular gene. Indeed, the signal peptide is the most variable region in the MBL sequence. While many changes in the signal peptide sequence are likely due to “neutral” genetic drift, some may be associated with co-evolution with the host translocation system. Developing our understanding of the universal rules for the relationship between signal peptide sequences and protein translocation should be a priority if we are to better characterize these important systems.

HGT is the primary mechanism underlying the dissemination of antibiotic resistance genes to different bacterial hosts. Needless to say, the host-compatibility of associated genetic elements such as the transcription and translation initiation sequences, and the origin of replication (in the case of the plasmids) have a significant impact on the outcome of HGT. However, our results suggest that coding genetic variation can itself play a significant role in a gene’s dissemination. It has been shown that acquired MBLs exhibit some bias with respect to the bacterial hosts that harbour them. For example, SPM-type MBLs have been noted for their strong association with *P. aeruginosa* (51), whereas NDM-type MBLs are predominantly found in *K. pneumoniae* and *E. coli* (**Table 1**) (52). Our study’s results are broadly consistent with these clinical observations, *i.e*., SPM-1 confers relatively high resistance in *P. aeuginosa*, but much lower resistance in *E. coli* and *K. pneumonia.* Moreover, a chromosomally encoded MBL, CcrA from *Bacteroides fragilis* appears to be toxic when expressed in *E. coli* and *P. aeuginosa* despite conferring substantial resistance in *K. pneumonia.* Beyond such examples, however, we did not observe any particular trend that differentiates acquirable MBLs from non-mobile MBLs. A more comprehensive survey using diverse clinical strains and the integration with other functional genetic elements, such as the native ribosome-binding site and promoter, may be needed to detect a more pronounced tendency (53). Additionally, there may be other specific features of MBLs that could be associated with high antibiotic resistance of pathogens in the human body. For example, recent studies demonstrated that NDM-1 variants possess the unique quality of being active in a partially zinc-depleted environment (54).

Lastly, our observations emphasize the importance of establishing comprehensive methodologies and protocols when studying newly isolated antibiotic resistance genes and variants. Antibiotic resistance variants continue to be isolated in clinical environments in increasing numbers, and in general, there has been more emphasis placed on determining their enzymatic efficiencies, with less concern paid to the detectable expression (55). We suggest that more effort is warranted for developing universal procedures and protocols. For example, the use of a universal expression vector for multiple host expression ability and representative clinically related model bacterial hosts to assess MBL variants *in situ* would allow for a more complete assessment of new clinical threats and facilitate the identification of hidden molecular properties that may be causing higher antibiotic resistance.

## Methods and Materials

### Construction of sequence similarity networks

A representative sequence similarity network (56,57) for the MBL B1 family was created by performing a protein BLAST on VIM-2 and then downloading all sequences with e-values of 10e^-15^ and lower on 23 March 2017. This cutoff was enough to include the adjacent B2 family seen in the full representative network, but limited other members of the superfamily. The sequences were filtered by length using the Galaxy Bioinformatics Suite (175 to 425 amino acids) before culling duplicate sequences with CD-HIT by using an identity threshold of 100%. The remaining 1199 sequences were added to 25 representative B1 and B2 MBL sequences (CphA, ImiS, Sfh-1, BlaB, EBR-1, SFB-1, SLB-1, GIM-1, SIM-1, KHM-1, DIM-1, PEDO-3, MUS-1, MUS-2, JOHN-1, CGB-1, TUS-1, BcII, CcrA, IND-1, NDM-1, VIM-1, VIM-2, IMP-1, and SPM-1) before an all versus all protein BLAST (National Center for Biotechnology Information, version 2.2.28+) with a threshold cutoff of 10e^-55^. The network was visualized with an organic layout in Cytoscape.

### Cloning and expression of MBLs in E. coli, P. aeruginosa, and K. pneumoniae

The eight MBL genes were synthesized (BioBasic) and subcloned in a broad-host-range vector, pBBR1MCS-2, along with the P_BAD_ promoter, which is inducible in the presence of arabinose (5.9-vector). The plasmids were transformed into three bacterial strains: *E. coli 10G* (chemical transformation), *P. aeruginosa* PA01 (electroporation) (58), and *K. pneumoniae* ATCC13883 (electroporation) (59).

### Determination of minimum inhibitory concentration (MIC) values

To determine the MIC for each <u>β</u>-lactam antibiotic, single colonies of *E. coli* 10G, *P. aeruginosa* PA01, and *K. pneumoniae* ATCC 13883 that were transformed with the MBL plasmids were picked, grown in quadruplicate overnight at 30°C in a 96-well plate in 500 μL of LB media with 1% glucose to suppress expression and 40 μg/mL of kanamycin for selective resistance. The overnight culture was used at a 1:40 dilution to start a new 200 μL culture of LB media supplemented with 40 μg/mL of kanamycin and 0.02% arabinose to induce expression. The cells were grown for 6 hours at 37°C before being transferred with replicator pins onto a series of LB agar plates containing two-fold increases in the concentration of antibiotic and 0.02% arabinose and grown overnight at 37°C (the range of concentrations screened for each antibiotic was as follows: cephalosporins, 0.032 μg/mL to 4096 μg/mL; carbapenams, 0.016 μg/mL to 64 μg/mL; penams, 2 μg/mL to 32768 μg/mL). The MICs were determined at the concentration of antibiotics by which no growth was observed at least three of the four replicates.

### Purification of Strep-tagged MBLs

All MBL variants were cloned into a pET-26(b) vector without their signal peptide and with a C-terminal Strep-tag (GNSGSAWSHPQFEK). Each enzyme was expressed in *E. coli* BL21 (DE3) cells in TB auto-induction media (EMD Millipore) supplemented with 1% (w/v) glycerol, 200 μM ZnCl_2_, and 40 μg/mL kanamycin. 200 mL cultures were inoculated with 5 mL of overnight culture (LB media, 40 μg/mL of kanamycin) and incubated at 30°C for 6 hours before further incubation at 18°C for 10 hours. Cells were harvested by centrifugation at 3200 × g and pellets were frozen at -80°C overnight. Cell pellets were resuspended in the lysis buffer containing 50% B-PER protein extraction reagent (Thermo Scientific) in Buffer A (50 mM Tris–HCl (pH 7.5), 100 mM NaCl and 200 μM ZnCl2) and 100 μg/mL of lysozyme, and incubated on ice for 1 hour. The cell lysates were centrifuged at 25,000 × g for 30 minutes at 4°C and the Strep-tag fusion proteins were purified from the clarified lysate according to the manufacturer’s instruction with Strep-tactin resin (IBA Lifesciences). The purified protein solution was desalted using Econo-Pac 10DG Column (Bio-Rad) and eluted in 4 mL of Buffer H (20 mM HEPES pH 7.5, 100 mM NaCl_2_, 200 μM ZnCl2). The concentration of each protein was determined by spectrophotometer. The A_280_ was measured for each sample with the following extinction coefficients: BcII, 34,950 M^-1^cm^-1^; IND-1, 40,910 M^-1^cm^-1^; IMP-1, 50,420 M^-1^cm^-1^; CcrA, 46,410 M^-1^cm^-1^; VIM-1, 33,920 M^-1^cm^-1^; VIM-2, 35,410 M^-1^cm^-1^; NDM-1, 33,460; SPM-1, 36,440 M^-1^cm^-1^.

### Enzyme assays to determine kinetic parameters

The catalytic ability of each MBL enzyme was measured for seven β-lactam substrates (CENTA, cefotaxime, ceftazidime, meropenem, imipenem, ampicillin, and benzylpenicillin) in Buffer H supplemented with 0.2% Triton X-100. The rate of hydrolysis of CENTA was determined by measuring changes in the absorbance at 405 nm (the extinction coefficient of CENTA at 405nm is 6400 M^-1^cm^-1^). The rate of hydrolysis of the β-lactam ring in the six antibiotics was determined by measuring changes in absorbance at the following wavelengths with these reported extinction coefficients: cefotaxime, 260 nm, -7500 M^-1^cm^-1^; ceftazidime, 260 nm, -9000 M^-1^cm^-1^; meropenem, 300 nm, -6500 M^-1^cm^-1^; imipenem, 300 nm, -9000 M^-1^cm^-1^; ampicillin 235 nm, -820 M^-1^cm^-1^; and benzylpenicillin, 235 nm, -775 M^-1^cm^-1^. The initial rates of reaction were measured in triplicate over the range of substrate concentrations (1 μM to 400 μM for CENTA, cefotaxime, ceftazidime, meropenem and imipenem, and 25 μM to 2000 μM for ampicillin and benzylpenicillin). The rates were used to determine the kinetic constants for each enzyme-substrate pair by fitting the data with the Michaelis-Menten equation using KaleidaGraph (Synergy). For those pairs where substrate saturation of the enzyme was not possible, the linear portion of the Michaelis-Menten plot was used to determine the *k*_cat_/*K*_M_ values.

### Cellular fractionation and the β-lactamase activity measurement in the cell lysate

To quantify the expression of the MBLs in the periplasm, cytoplasm, and whole cell fractions, single colonies from *E. coli* 10G, *P. aeruginosa* PA01, and *K. pneumoniae* ATCC 13883 transformed with the MBL plasmids were picked, grown in quadruplicate overnight at 30°C in a 96-well plate in 200 μL of LB media with 1% glucose to suppress expression and 40 μg/mL of kanamycin for selective resistance. A 1:40 dilution was used to start a 500 μL LB media expression culture that was grown for 6 hours at 37°C with 0.02% arabinose for induction and 40 μg/mL of kanamycin. The cells were collected by centrifugation at 3200 × g for 10 minutes. The periplasmic fractions were isolated using the osmotic shock protocol. The pellets were suspended in OS1 buffer (30 mM Tris-HCl, pH 7.1, 20% sucrose, and 1mM phenylmethylsulfonyl fluoride) and incubated for 30 minutes at room temperature before centrifugation (3200 × g for 10 minutes). The pellets were then resuspended in OS2A buffer (ice-cold 0.5 mM MgCl_2_) and incubated on ice for 5 minutes before centrifugation (3200 × g for 10 minutes). 100 μL of the supernatant was collected and mixed with 100 μL OS2B (40 mM HEPES, pH 7.5, 200 mM NaCl, 400 μM ZnCl_2_, and 0.4% Triton). The initial rate of β-lactamase activity was measured by mixing the isolated periplasm fraction and 50 μM of CENTA at a 1:10 ratio in Buffer H supplemented with 0.2% Triton X-100. The remaining pellets were then frozen overnight at -20°C. To obtain the cytoplasmic fraction, the frozen pellets were resuspended in 200 μL of lysis buffer (20 mM HEPES pH 7.5, 100 mM NaCl, 200 μM ZnCl2, 0.2% Triton, 200 ug/mL Lysozyme, 1 U Benzonase (Millipore), and 0.25 mM MgCl_2_), and incubated at room temperature for 1 hour. After centrifugation (3200 × g for 10 minutes), the supernatant was removed and β-lactamase activity was measured to determine the level of enzymes in the cytoplasm. The β-lactamase activity of the whole cell was prepared in the same way as the cytoplasmic fraction without the initial isolation of the periplasmic fraction.

### Calculation of the number of functional MBLs per fraction

The initial rate of reaction with CENTA for the fractional or whole cell lysates was used in conjunction with the initial rate measured with purified enzyme to determine the relative amount of enzyme in each fraction: [*E*] = ((*A*_lysate_ ÷ *A*_purified_) × [*E*_purified_] × *V* x *N*_A_) ÷ (OD_600_ × 1×10^9^ CFU/OD_600_), where *A*_lysate_ denotes the rate of CENTA hydrolysis in the lysate, *A*_purified_ denotes the rate of CENTA hydrolysis of purified enzyme, [*E*_purified_] denotes the concentration of purified enzyme, *V* denotes the culture volume, *N*_A_ denotes Avogadro’s Constant, and OD_600_ denotes the absorbance of the cultures at 600 nm.

### Determination of MBL gene expression

After induction of the *E. coli* harbouring each MBL as described above, plasmid DNA was isolated from 500 μL of culture with a QIAprep Miniprep Kit (Qiagen). For isolation of RNA, 500 μL of the induced culture was mixed with 500 μL of RNAprotect Bacteria Reagent (Qiagen) and incubated at room temperature for 5 minutes before RNA extraction as per the manufacturer’s instructions with the RNeasy Mini Kit (Qiagen). To ensure removal of contaminating DNA, the samples were processed with a DNA-Free kit (Ambion). cDNA was then prepared using the QuantiTect Reverse Transcription Kit (Qiagen).

Quantitative PCR was performed on a 7500 Fast Real-Time PCR System (Applied Biosystems) with the following cycling conditions: 20 seconds at 95°C, followed by 40 cycles of 95°C for 3 seconds and 58°C for 30 seconds. The primers used for the DNA quantification were designed to detect the pBBR1-pBAD vector: F-CCAACAGCGATTCGTCCTGG, R-AGCCAGAAGACACTTTCCAAGC. The forward primers used for the RNA quantification were designed for each MBL: BcII, F-GATTTAGGAAACGTTGCGGATGC; IND-1, F-CAATGTATTGGATGGTGGCTGTC; CcrA, F-GCATGGCCGAAAACTCTCG; NDM-1, F-CTCGGCAATCTCGGTGATGC; VIM-1, F-GGAAGCAGAGGTCGTCATTCC; VIM-2, F-GGAAGCACAGTTCGTCATTCC; IMP-1, F-TAGAAGCTTGGCCAAAGTCCG; and SPM-1, F-AACTTGGTTATCTGGGAGATGCC. The reverse primer was the same for all eight MBLs: R-GCAACGCAATTAATGTGAGTTAGC. The primers for the reference gene, histidyl-RNA synthetase (*hisS*), were: F-GCTCCGGCATTAGGTGATTA and R-TCAAGCAGTTTGCACAGACC. Normalized expression units for each MBL gene were calculated using the ΔΔCt method relative to *hisS*, whereas the absolute value of the number of DNA copy number was determined with known standards of the vector, pBBR1-pBAD, and normalized by the OD_600_ of the original culture.

### Replacement of MBL signal peptides with the PelB leader sequence

The native signal peptides for each MBL were replaced in the pBBR1-pBAD vector with the PelB leader sequence (MGKYLLPTAAAGLLLLAAQPAMAMDSG) using Golden Gate assembly cloning with the Type IIS restriction enzyme, *Bsa*I.

## Acknowledgements

We thank the members of the Tokuriki lab for comments on the manuscript. Canadian Institute of Health Research (CIHR) Foundation Grant to N.T.. N.T. is a CIHR new investigator and a Michael Smith Foundation of Health Research (MSFHR) career investigator.

## Author contributions

R.D.S. and N.T. conceived and designed this study. R.D.S performed all experiments R.D.S. and N.T. wrote the paper.

